# An Integrative Analysis of the Age-Associated Genomic, Transcriptomic and Epigenetic Landscape across Cancers

**DOI:** 10.1101/2020.08.25.266403

**Authors:** Kasit Chatsirisupachai, Tom Lesluyes, Luminita Paraoan, Peter Van Loo, João Pedro de Magalhães

## Abstract

Age is the most important risk factor for cancer, as cancer incidence and mortality increase with age. However, how molecular alterations in tumours differ among patients of different age remains largely unexplored. Here, using data from The Cancer Genome Atlas, we comprehensively characterised genomic, transcriptomic and epigenetic alterations in relation to patients’ age across cancer types. We showed that tumours from older patients present an overall increase in genomic instability, somatic copy-number alterations (SCNAs) and somatic mutations. Age-associated SCNAs and mutations were identified in several cancer-driver genes across different cancer types. The largest age-related genomic differences were found in gliomas and endometrial cancer. We identified age-related global transcriptomic changes and demonstrated that these genes are controlled by age-associated DNA methylation changes. This study provides a comprehensive view of age-associated alterations in cancer and underscores age as an important factor to consider in cancer research and clinical practice.

## Introduction

Age is the biggest risk factor for cancer, as cancer incident and mortality rates increase exponentially with age in most cancer types^1^. However, the relationship between ageing and molecular determinants of cancer remains to be characterised. Cancer arises through the interplay between somatic mutations and selection, in a Darwinian-like process^2,3^. Thus, apart from the mutation accumulation with age^4–6^, microenvironment changes during ageing could also play a role in carcinogenesis^2,7,8^. We therefore hypothesise that, due to the differences in selective pressures from tissue environmental changes with age, tumours arise from patients across different ages might harbour different molecular landscapes; consequently, some molecular changes might be more or less common in older or younger patients.

Recently, several studies have investigated the molecular differences in the cancer genome in relation to clinical factors, including gender^9,10^ and race^11,12^. These studies demonstrated gender- and race-specific biomarkers, actionable target genes and provided clues to understanding the biology behind the disparities in cancer incidence, aggressiveness and treatment outcome across patients from different backgrounds. Although the genomic alterations in childhood cancers and the differences with adult cancers have been systematically characterised^13,14^, the age-related genomic landscape across adult cancers remains elusive. Specific age-associated molecular landscapes have been reported in the cancer genome of several cancer types, for example, glioblastoma^15^, prostate cancer^16^ and breast cancer^17^. However, these studies focused mainly on a single cancer type and only on some molecular data types.

Here, using data from The Cancer Genome Atlas (TCGA), we systematically investigated age-related differences in genomic instability, somatic copy number alterations (SCNAs), somatic mutations, pathway alterations, gene expression, and DNA methylation landscape across various cancer types. We show that, in general, genomic instability and mutations frequency increase with age. We identify several age-associated genomic alterations in cancers, particularly in low-grade glioma and endometrial carcinoma. Moreover, we also demonstrate that age-related gene expression changes are controlled by age-related DNA methylation changes and that these changes are linked to numerous biological processes.

## Results

### Association between age and genomic instability, loss of heterozygosity, and whole-genome duplication

To gain insight into the role of patient age into the somatic genetic profile of tumours, we evaluated associations between patient age and genomic features of tumours in TCGA data (Table 1, Supplementary Table 1). Using multiple linear regression adjusting for gender, race, and cancer type, we found that genomic instability (GI) scores increase with age in pan-cancer data (adj. R-squared = 0.35, p-value = 5.98×10^−7^) (Fig. 1a). We next applied simple linear regression to investigate the relationship between GI scores and age for each cancer type. Cancer types with a significant association (adj. p-value < 0.05) were further adjusted for clinical variables. We found a significant positive association between age and GI score in seven cancer types (adj. p-value < 0.05) (Fig. 1b, Supplementary Fig. 1a and Supplementary Table 2). Cancer types with the strongest significant positive association were low-grade glioma, ovarian cancer, endometrial cancer, and sarcoma. This result indicates that the level of genomic instability increases with the age of cancer patients in several cancer types.

**Table 1.** Summary of TCGA cancer type and number of samples used in each analysis

| Cancer type | Abbreviation | GI, LOH, WGD | SCNAs | Mutations (hypermutated tumours removed) | Pathway alterations | Gene expression | DNA methylation |
| --- | --- | --- | --- | --- | --- | --- | --- |
| Adrenocortical carcinoma | ACC | 89 | 89 | 89 (88) | 76 | 77 | 78 |
| Bladder Urothelial Carcinoma | BLCA | 370 | 369 | 369 (364) | 361 | 366 | 370 |
| Breast invasive carcinoma | BRCA | 1015 | 1011 | 954 (946) | 922 | 1011 | 719 |
| Cervical squamous cell carcinoma and endocervical adenocarcinoma | CESC | 287 | 287 | 271 (263) | 264 | 284 | 287 |
| Cholangiocarcinoma | CHOL | 35 | 35 | 35 (35) | 35 | 35 | 35 |
| Colon adenocarcinoma | COAD | 411 | 411 | 374 (319) | 323 | 410 | 278 |
| Lymphoid Neoplasm Diffuse Large B-cell Lymphoma | DLBC | 42 | 42 | 32 (32) | 32 | 42 | 42 |
| Oesophageal carcinoma | ESCA | 176 | 176 | 176 (174) | 165 | 175 | 176 |
| Glioblastoma multiforme | GBM | 489 | 489 | 356 (354) | 116 | 137 | 259 |
| Head and Neck squamous cell carcinoma | HNSC | 489 | 489 | 472 (469) | 459 | 481 | 489 |
| Kidney Chromophobe | KICH | 66 | 66 | 66 (66) | 65 | 66 | 66 |
| Kidney renal clear cell carcinoma | KIRC | 496 | 496 | 343 (343) | 331 | 493 | 296 |
| Kidney renal papillary cell carcinoma | KIRP | 228 | 228 | 222 (222) | 215 | 228 | 213 |
| Acute Myeloid Leukaemia | LAML | 126 | 121 | 55 (54) | 101 | 102 | 121 |
| Brain Lower Grade Glioma | LGG | 488 | 488 | 484 (484) | 482 | 488 | 488 |
| Liver hepatocellular carcinoma | LIHC | 355 | 355 | 342 (340) | 334 | 349 | 355 |
| Lung adenocarcinoma | LUAD | 460 | 460 | 456 (438) | 446 | 456 | 402 |
| Lung squamous cell carcinoma | LUSC | 460 | 460 | 444 (437) | 426 | 457 | 336 |
| Mesothelioma | MESO | 82 | 82 | 77 (77) | 77 | 82 | 82 |
| Ovarian serous cystadenocarcinoma | OV | 556 | 556 | 397 (395) | 173 | 288 | 545 |
| Pancreatic adenocarcinoma | PAAD | 133 | 133 | 130 (129) | 113 | 127 | 132 |
| Pheochromocytoma and Paraganglioma | PCPG | 165 | 157 | 165 (164) | 154 | 165 | 165 |
| Prostate adenocarcinoma | PRAD | 434 | 434 | 434 (432) | 425 | 434 | 434 |
| Rectum adenocarcinoma | READ | 152 | 152 | 132 (127) | 109 | 151 | 95 |
| Sarcoma | SARC | 229 | 229 | 213 (211) | 209 | 227 | 229 |
| Skin Cutaneous Melanoma | SKCM | 434 | 434 | 432 (340) | 332 | 433 | 434 |
| Stomach adenocarcinoma | STAD | 388 | 388 | 385 (345) | 340 | 365 | 341 |
| Testicular Germ Cell Tumours | TGCT | 129 | 129 | 124 (124) | 123 | 129 | 129 |
| Thyroid carcinoma | THCA | 260 | 260 | 249 (248) | 244 | 259 | 258 |
| Thymoma | THYM | 76 | 76 | 76 (76) | 73 | 73 | 76 |
| Uterine Corpus Endometrial Carcinoma | UCEC | 434 | 434 | 421 (366) | 406 | 432 | 360 |
| Uterine Carcinosarcoma | UCS | 52 | 52 | 52 (51) | 52 | 52 | 52 |
| Uveal Melanoma | UVM | 72 | 72 | 72 (72) | 72 | 72 | 72 |
| Total |  | 9678 | 9660 | 8899 (8585) | 8055 | 8946 | 8414 |

**Fig. 1.**
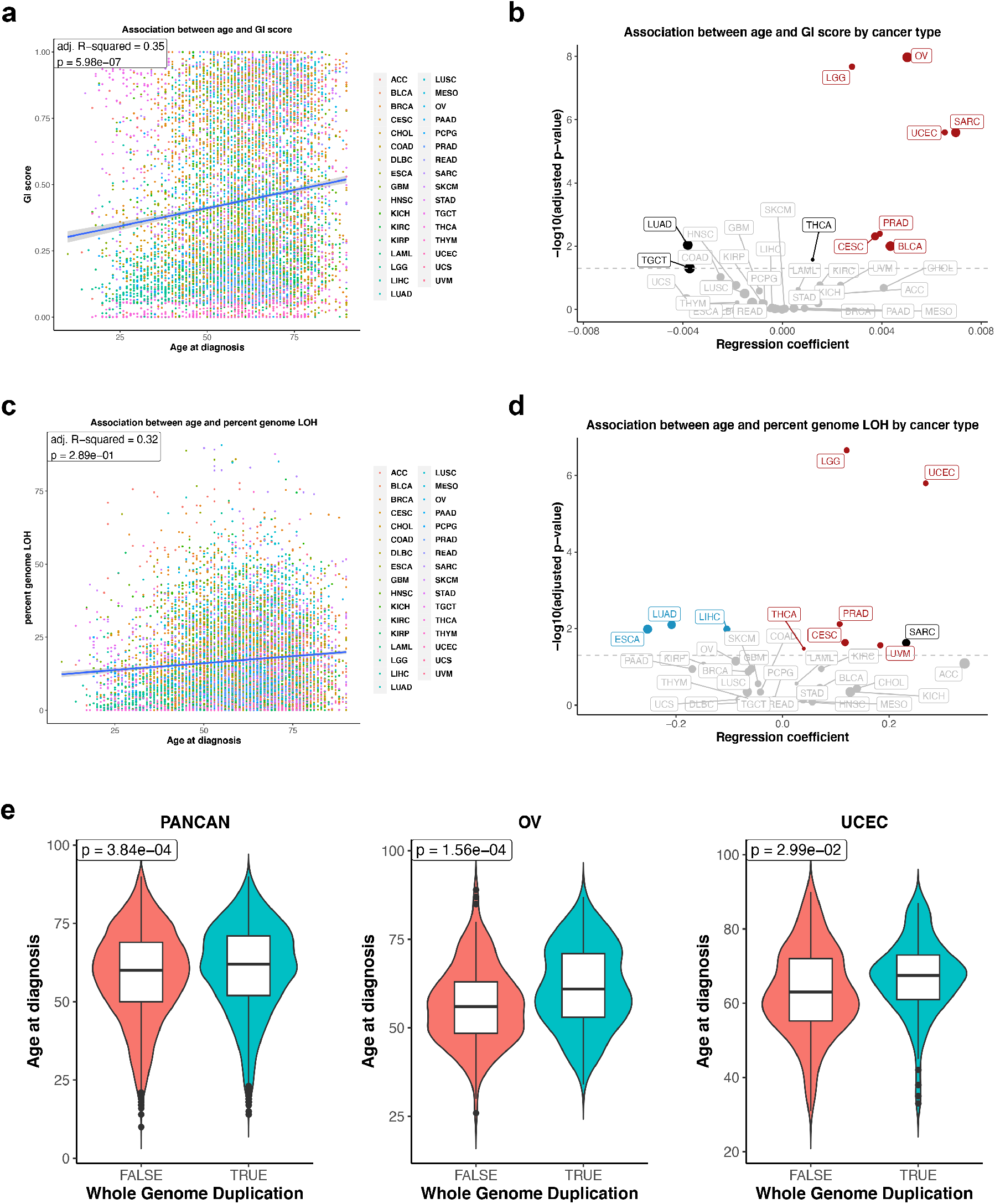
Association between cancer patients’ age and genomic instability (GI) score, percent genomic loss-of-heterozygosity (LOH) and whole-genome duplication events (WGD). **a** Association between age and pan-cancer GI score. Dots are coloured by cancer type. Multiple linear regression R-squared and p-value are shown in the figure. **b** Association between age and cancer type-specific GI score. Linear regression coefficients and significant values are shown in the figure. Cancers with a significant positive association between age and GI score after using multiple linear regression (adj. p-value < 0.05) are highlighted in red. Cancers with a significant association in simple linear regression but not significant after using multiple linear regression are showed in black. The grey line indicates adj. p-value = 0.05. Dot size is proportional to median GI score. **c** Association between age and pan-cancer percent genomic LOH. Dots are coloured by cancer type. Multiple linear regression R-squared and p-value are shown in the figure. **d** Association between age and cancer type-specific percent genomic LOH. Simple linear regression results are shown. Cancers with a significant positive and negative association between age and percent genomic LOH after using multiple linear regression are highlighted in red and blue, respectively. Cancer with a significant association in simple linear regression but not significant after using multiple linear regression is showed in black. The grey line indicates adj. p-value = 0.05. Dot size is proportional to median percent genomic LOH. **e** Association between age and WGD events in pan-cancer, OV, and UCEC. Multiple logistic regression p-values were indicated in the figure.

The genomic loss of heterozygosity (LOH) refers to the irreversible loss of one parental allele, causing an allelic imbalance, and priming the cell for another defect at the other remaining allele of the respective genes^18^. To investigate whether there is an association between patients’ age and LOH, we quantified percent genomic LOH. By using simple linear regression, we found a significant positive association between age and pan-cancer percent genomic LOH (p-value = 1.20 × 10^−21^). However, this association was no longer significant in a multiple linear regression analysis (adj. R-squared = 0.32, p-value = 0.289) (Fig. 1c). Thus, it is likely that this association might be cancer type-specific. We then performed a linear regression between age and percent genomic LOH for each cancer type. Six cancer types showed a positive association between age and percent genomic LOH (adj. p-value < 0.05) (Fig. 1d, Supplementary Fig. 1b, and Supplementary Table 3). The strongest positive associations were found in low-grade glioma and endometrial cancer (adj. p-value < 0.05), corroborate with the increase in GI score with age. On the other hand, lung adenocarcinoma, oesophageal and liver cancer demonstrated a negative correlation between percent genomic LOH and age (adj. p-value < 0.05).

Whole-genome duplication (WGD) is important in increasing the adaptive potential of the tumour and has been linked with a poor prognosis^19–21^. We investigated the relationship between age and WGD using logistic regression. For the pan-cancer analysis, we found an increase in the probability that WGD occurs with age, using multiple logistic regression accounting for gender, race, and cancer type (odds ratio per year (OR) = 1.0066, 95% confidence interval (CI) = 1.0030-1.0103, p-value = 3.84 × 10^−4^) (Fig. 1e). For the cancer-specific analysis, a significant positive association was found in ovarian and endometrial cancer (adj. p-value < 0.05, OR = 1.0320 and 1.0248, 95%CI = 1.0151-1.0496 and 1.0024-1.0483, respectively) (Fig. 1e and Supplementary Table 4), indicating that tumours from older patients are more likely to have doubled their genome. Taken together, the findings indicate that tumours from patients with an increased age tend to harbour a more unstable genome and a higher level of LOH in several cancer types. Notably, the strongest association between age and an increase in genome instability, LOH, and WGD was evident in endometrial cancer, suggesting the potential disparities in cancer genome landscape with age in this cancer type.

### Age-associated somatic copy-number alterations

We used GISTIC2.0 to identify recurrently altered focal- and arm-level SCNAs^22^. We calculated the SCNA score, as a representation of the level of SCNA occurring in a tumour^12,23^. For each tumour, the SCNA score was calculated at three different levels: focal-, arm- and chromosome-level, and the overall score calculated from the sum of all three levels. We used simple linear regression to identify the association between age and overall SCNA scores. Cancer types that displayed a significant association were further adjusted for clinical variables. Consistent with the GI score results described above, the strongest positive association between age and overall SCNA scores was found in low-grade glioma, ovarian and endometrial cancers. Other cancer types for which a positive association between age and overall SCNA score was observed were thyroid cancer and clear cell renal cell carcinoma (adj. p-value < 0.05). On the other hand, lung adenocarcinoma is the only cancer type exhibiting a negative association between overall SCNA score and age (Fig. 2a, Supplementary Fig. 2a, and Supplementary Table 5). The different SCNA classes (focal- and chromosome/arm-level) may arise through different biological mechanisms^12,21^, therefore we separately analysed the association between age and focal- and chromosome/arm-level SCNA scores. Most cancers that showed a significant relationship between age and overall SCNA score also had an association between age and both chromosome/arm-level and focal-level SCNA scores (Fig. 2b-c, Supplementary Fig. 2b-c, and Supplementary Table 5). The only exception was in cervical cancer, with a significant association between age and chromosome/arm-level but not with focal-level and overall SCNA scores.

**Fig. 2.**
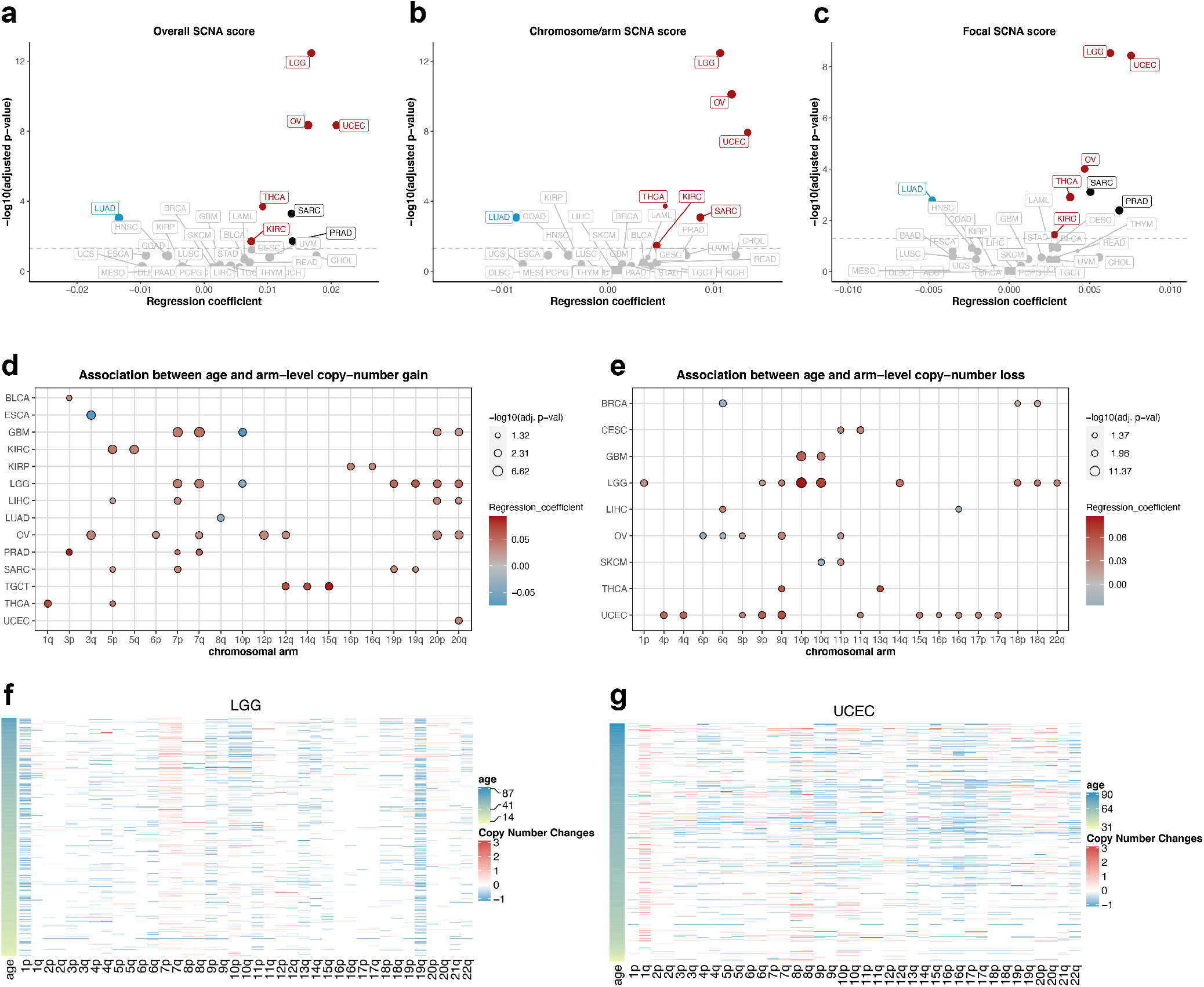
Association between cancer patients’ age and somatic copy-number alterations (SCNAs). Volcano plot representing the association between age and **(a)** overall, **(b)** focal-level and **(c)** chromosome/arm-level SCNA scores. Linear regression coefficients and significant values are shown. Cancers with a significant positive and negative association between age and SCNA score after using multiple linear regression (adj. p-value < 0.05) are highlighted in red and blue, respectively. Cancers with a significant association in simple linear regression but not significant after using multiple linear regression are showed in black. The grey line indicates adj. p-value = 0.05. Dot size is proportional to median SCNA score. **d, e** The left and right dot plots show the association between age and arm-level copy-number gains and copy-number deletions. Circle size corresponds to the significant level, red and blue represent positive and negative associations, respectively. **f, g** Heatmaps represent recurrently gain and deletion arms in LGG and UCEC, respectively. Samples are sorted by age. Colours represent copy-number changes from GISTIC2.0, blue denotes deletion and red corresponds to gain.

We next identified the chromosomal arms that tend to be gained and lost more often with age, for 25 cancer types with sufficient samples (at least 100 tumours, Table 1). We conducted the logistic regression on the significant recurrently gained and lost arms that were identified by GISTIC2.0 for each cancer type. The significant association between age and chromosomal arm gains and losses are shown in Fig. Fig. 2d, e, respectively (adj. p-value < 0.05) (Supplementary Fig. 3, Supplementary Table 6). The gain of chromosome 7p, 7q, 20p, and 20q significantly increased with age in several cancer types including two types of gliomas, low-grade glioma and glioblastoma. On the other hand, the gain of chromosome 10p decreased with increased age in gliomas (Fig. 2d and 2f). For the arm losses, there was an increased occurrence of loss in 11 arms with advanced age in endometrial cancer (Fig. 2e and 2g), consistent with a higher genomic instability and LOH with age in this cancer type. Low-grade glioma and ovarian cancer, two other cancer types for which we found the highest significant association between age and SCNA scores, also exhibited a significant increase or decrease in losses with age in multiple arms (Fig. 2e-f, Supplementary Fig. 3). We also observed that the losses of chromosome 10p and 10q increased with age in gliomas. Recurrent losses of chromosome 10 together with the gain of chromosome 7 are important features in IDH-wild-type (IDH-WT) gliomas^24^. This type of gliomas was more common in older patients, whereas IDH-mutated gliomas were predominantly found in younger patients.

We further examined age-associated recurrent focal-level SCNAs. Applying a similar logistic regression, we identified recurrent focal SCNAs associated with the age of the patients for each cancer type. In total, we found 113 significant age-associated regions, including 67 gain regions across 10 cancer types and 46 loss regions across 9 cancer types (adj. p-value < 0.05) (Fig. 3a, Supplementary Table 7). In accordance with the arm-level result, the highest number of significant regions was found in endometrial cancer (23 gain and 25 loss regions), followed by ovarian cancer (13 gain 2 loss regions) and low-grade glioma (9 gain and 5 loss regions) (Fig. 3b-c, Supplementary Fig. 4).

**Fig. 3.**
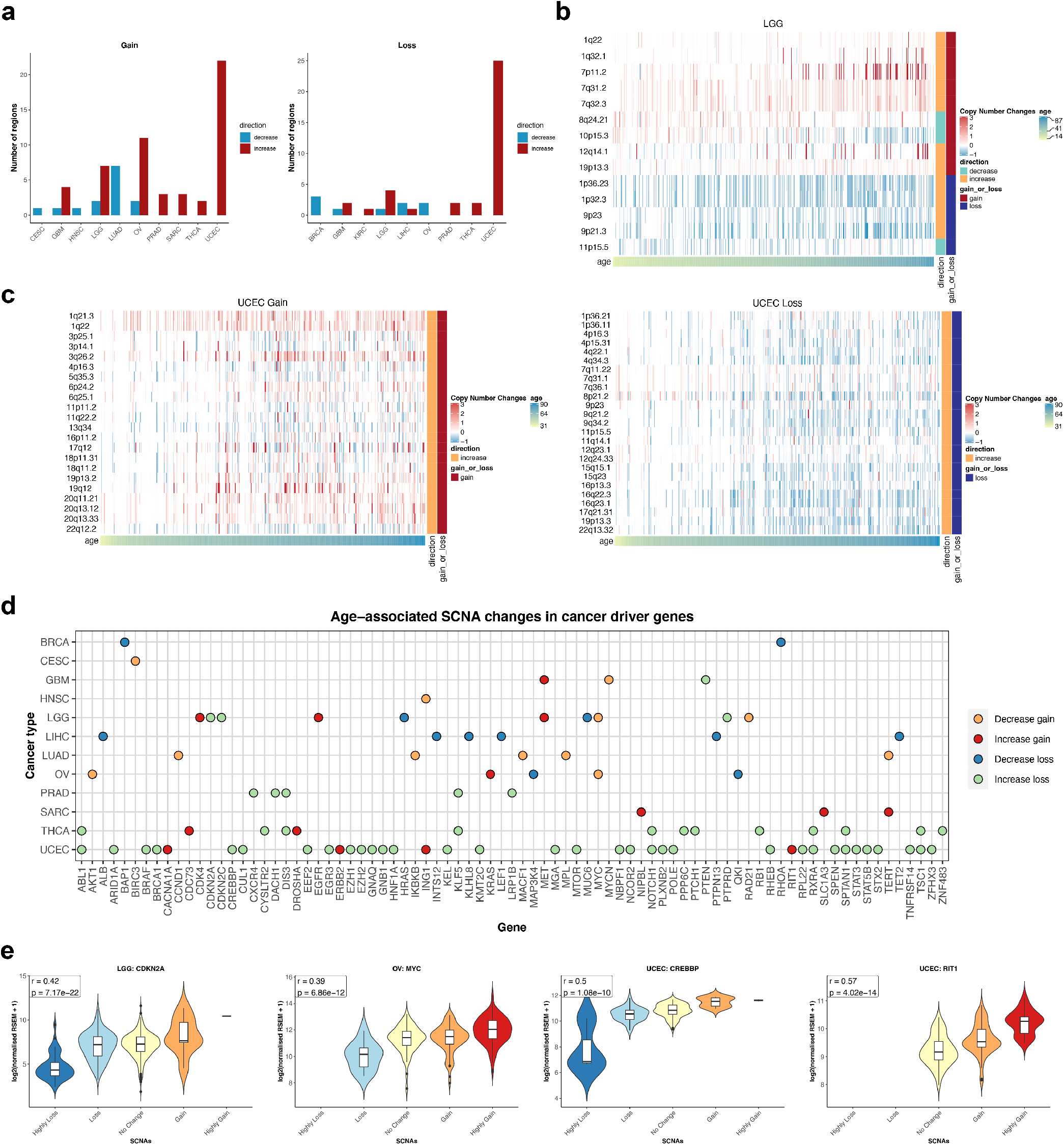
Association between cancer patients’ age and focal-level SCNAs. **a** Number of gained and deleted focal regions that showed a significant association with age per cancer type (multiple logistic regression, adj. p-value < 0.05). Heatmap showing age-associated focal-level SCNAs in **(b)** LGG and **(c)** UCEC. Samples are sorted by age. Colours represent copy-number changes from GISTIC2.0, blue denotes deletion and red corresponds to gain. The gain_or_loss legend demonstrates that the region is recurrently gained or deleted. The direction legend shows whether the gain/deletion of the region increases or decreases with age. **d** Age-associated SCNA changes in cancer driver genes. Cancer driver genes located in the age-associated focal regions are plotted by cancer type. Colours of the dot represent the condition of the focal region where the gene located in as follows: blue - decrease deletion; green - increase deletion; yellow - decrease gain; and red - increase gain with age. **e** The effect of copy-number changes on gene expression of *CDKN2A* in LGG, *MYC* in OV, *CREBBP* and *RIT1* in UCEC. These are examples of genes with age-associated changes in SCNAs. Violin plots show the log2(normalized expression + 1) of samples grouped by their SCNA status. Pearson correlation coefficient r and p-value are shown in the figures.

To further investigate the impact of these SCNAs, we studied the correlation between the SCNA level and gene expression for tumours that have both types of data using Pearson correlation. In total, 81 genes in the list of previously identified cancer driver genes (Supplementary Table 8) were presented in at least one significant age-associated focal region in at least one cancer type and showed a significant correlation between SCNAs and gene expression (adj. p-value < 0.05) (Fig. 3d). For example, regions showing an increased gain with age in endometrial cancer included 1q22, where the gene *RIT1* is located in (OR = 1.0355, 95%CI = 1.0151-1.0571, adj. p-value = 0.0018) (Fig. 3c, e). The Ras-related GTPases *RIT1* gene has been reported to be highly amplified and correlated with poor survival in endometrial cancer^25^. Therefore, an increase in the gain of the *RIT1* gene with age might relate to a poor prognosis in older patients. The 16p13.3 loss increased in older endometrial cancer patients (OR = 1.0335, 95%CI = 1.0048-1.0640, adj. p-value = 0.0328). This region contains the p53 coactivator gene *CREBBP*. The gain of 8q24.21 harbouring the oncogene *MYC* decreased with patient age in low-grade glioma (OR = 0.9737, 95%CI = 0.9541-0.9927, adj. p-value = 0.0128) and ovarian cancer (OR = 0.9729, 95%CI = 0.9553-0.9904, adj. p-value = 0.0063) (Fig. 3d, e). In addition, in low-grade glioma, we found an increase in 9p21.3 loss with age (OR = 1.0332, 95%CI = 1.0174-1.0496, adj. p-value = 0.00017). This region contains the cell cycle-regulator genes *CNKN2A* and *CDKN2B* (Fig. 3b, d, e). The full list of age-associated focal regions across cancer types and the correlation between SCNA status and gene expression can be found in Supplementary Table 7. Taken together, our analysis demonstrates the association between age and SCNAs level across cancer types. We also identified age-associated arms and focal-regions, and these regions harboured several cancer-driver genes. Our results suggest a possible contribution of different SCNA events in cancer initiation and progression of patients with different ages.

### Age-associated somatic mutations in cancer

The increase in the mutational burden with age is well-established^4–6^. This age-related mutation accumulation is largely explained by a clock-like mutational process, the spontaneous deamination of 5-methylcytosine to thymine^5^. As expected, we confirmed the correlation between age and mutation load (somatic non-silent SNVs and indels) in the pan-cancer cohort using multiple linear regression adjusting for gender, race, and cancer type (adj. R-squared = 0.53, p-value = 1.41 × 10^−37^) (Supplementary Fig. 5a). For cancer-specific analysis, 18 cancer types exhibited a significant relationship between age and mutation load using linear regression (adj. p-value < 0.05) (Supplementary Fig. 5, Supplementary Table 9). Only endometrial cancer showed a negative correlation between mutational burden and age. We observed a high proportion of hypermutated tumours (> 1,000 non-silent mutations per exome) from younger endometrial cancer patients. Thirteen out of 38 tumours (34%) from the younger patients (age ≤ 50) were hypermutated tumours, while there were only 42 hypermutated tumours from 383 tumours from older patients (11%) (Fisher’s exact, p-value = 0.0003) (Fig. 4a). Microsatellite instability (MSI) is a unique molecular alteration caused by defects in DNA mismatch repair^26,27^. The MSI-high (MSI-H) tumours occur as a subset of high mutational burden tumours^28^. We investigated whether high mutation loads in endometrial cancer from young patients were due to the presence of MSI-H tumours. Using multiple logistic regression, we found that MSI-H tumours were associated with younger endometrial cancer (OR = 0.9751, 95%CI = 0.9531-0.9971, p-value = 0.0264) (Fig. 4b). Another source of hypermutation in cancer is the defective DNA polymerase proofreading ability by mutations in polymerase ε (*POLE*) or polymerase δ (*POLD1*) genes^29,30^. We showed that mutations in *POLE* (OR = 0.9690, 95%CI = 0.9422-0.9959, p-value = 0.0243) and *POLD1* (OR = 0.9573, 95%CI = 0.9223-0.9925, p-value = 0.0177) were both more prevalent in younger endometrial cancer patients (Fig. 4c). Therefore, the negative correlation between age and mutation loads in endometrial cancer could be explained by the presence of hypermutated tumours in younger patients, which are associated with MSI-H and *POLE/POLD1* mutations. Previous studies on *POLE* and MSI-H subtypes in hypermutated endometrial tumours revealed that these subtypes associated with a better prognosis when compared with the copy-number high subtype^31–33^. Together with our SCNA results, younger UCEC patients are likely to associate with a *POLE* and MSI-H subtypes, high mutation rate and better survival, whilst tumours from older patients are characterized by high SCNAs and are generally associated with a worse prognosis. We extended the age and MSI-H analysis to other cancer types known to have a high prevalence of MSI-H tumours, including colon, rectal, and stomach cancers^26^. Only in stomach cancer we found an association between older age and the presence of MSI-H tumours (OR = 1.0392, 95%CI = 1.0091-1.0720, p-value = 0.01, Supplementary Fig. 6a). When we further examined the association between age and mutations in *POLE* and *POLD1* in other cancers apart from endometrial cancer, no significant association was observed (Supplementary Fig. 6b).

**Fig. 4.**
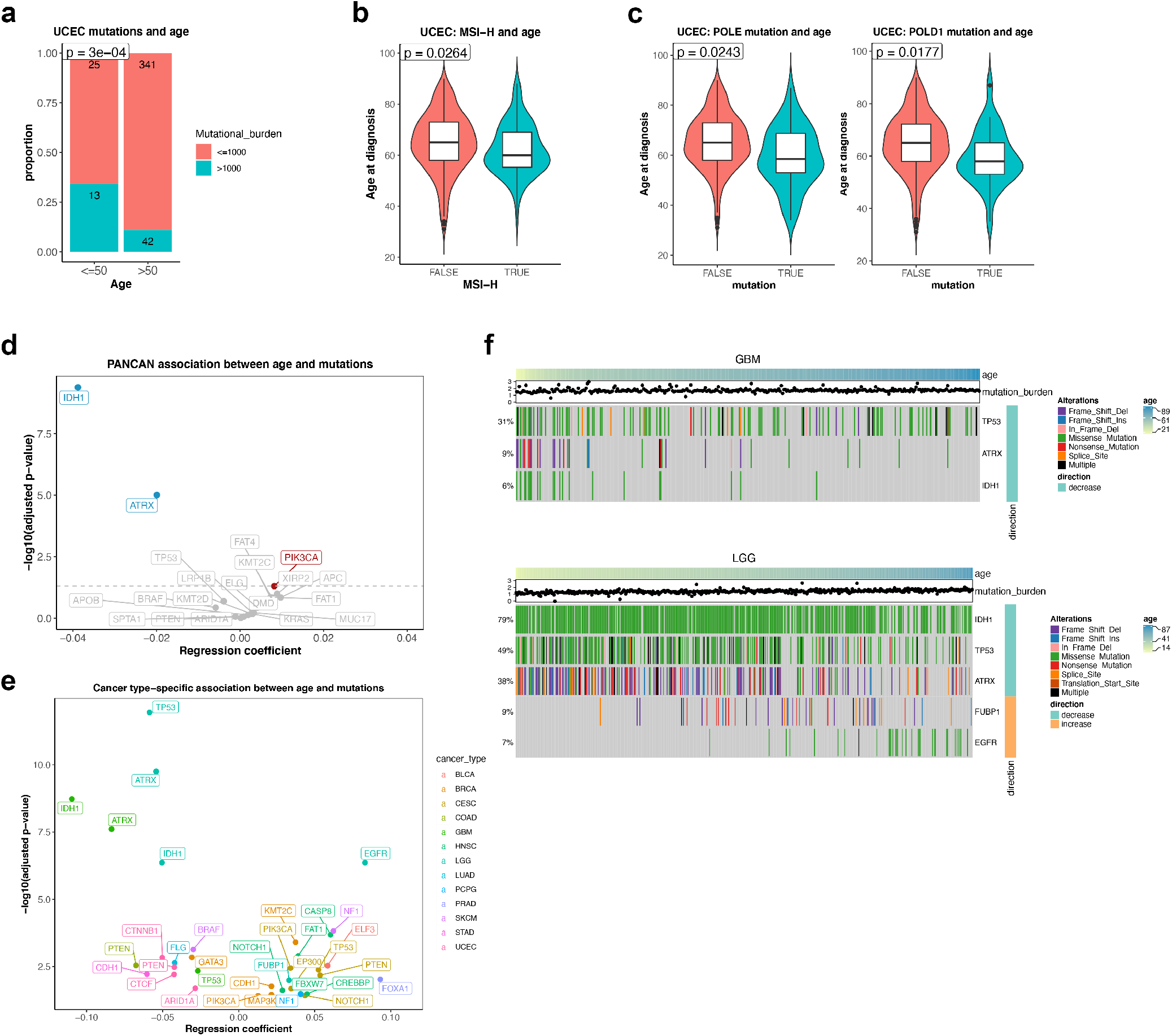
Association between cancer patients’ age and somatic mutations. **a** The proportion of hypermutated tumours (>1,000 mutations/exome) in young (age ≤ 50) and old (age > 50) UCEC. The statistical significant (p-value) was calculated using Fisher’s exact test. **b** The association between age and MSI-H in UCEC. The statistical significance was calculated from the multiple logistic regression adjusting for clinical variables. The p-value is shown in the figure. **c** The association between age and *POLE/POLD1* mutations in UCEC. The statistical significance (p-value) was calculated from the multiple logistic regression adjusting for clinical variables. **d** A pan-cancer association between age and mutations. Multiple logistic regression coefficient and significant values are shown. Genes with a significant positive and negative association between age and somatic mutations after using multiple logistic regression (adj. p-value < 0.05) are highlighted in red and blue, respectively. **e** Summary of the cancer type-specific association between age and mutations. Multiple logistic regression coefficient and significant values are shown. Only genes with a significant association (adj. p-value < 0.05) are shown in the figure. A colour code is provided to denote the cancer type where the association between age and gene mutation was found. **f** Heatmap showing age-associated mutations in GBM and LGG. Samples are sorted by age. Colours represent types of mutation. The right annotation legend indicates the direction of change, increase or decrease mutations with age. The mutational burden of samples is presented in the dot above the heatmap.

Although the increase in mutation load with age in cancer is well studied^4,28^, the bias of mutation in particular genes with age across cancer types is largely unclear. To better understand this, we conducted logistic regression to investigate genes that are more or less likely to be mutated with an increased age. To prevent the potential bias caused by hypermutated tumours, we restricted the analysis to samples with < 1,000 non-silent mutations per exome (Table 1). We first investigated the association between age and pan-cancer gene-level mutations. Using multiple logistic regression correcting for gender, race, and cancer type, mutations in *IDH1* (OR = 0.9619, 95%CI = 0.9510-0.9730, adj. p-value = 4.18 × 10^−10^) and *ATRX* (OR = 0.9803, 95%CI = 0.9724-0.9881, adj. p-value = 9.85×10^−6^) showed a negative association with age. On the other hand, mutations in *PIK3CA* were more common in older individuals (OR = 1.0082, 95%CI = 1.0022-1.0143, adj. p-value = 4.18×10^−10^) (Fig. 4d). We next identified genes in which mutations associated with age in a cancer-specific manner in 24 cancers with at least 100 samples (Table 1). Using logistic regression, we identified 35 mutations from 13 cancers that increased or decreased with the patients’ age (adj. p-value < 0.05) (Fig. 4e-f, Supplementary Fig. 7 and Supplementary Table 10). The most striking negative associations between mutations and age in low-grade glioma and glioblastoma were found in *IDH1* (OR = 0.9509 and 0.8962, 95%CI = 0.9328-0.9686 and 0.8598-0.9291, adj. p-value = 4.33×10^−7^ and 1.88×10^−9^, respectively), *ATRX* (OR = 0.9471 and 0.9120, 95%CI = 0.9310-0.9628 and 0.8913-0.9466, adj. p-value = 1.75×10^−10^ and 2.45×10^−8^, respectively), and *TP53* (OR = 0.9431 and 0.9736, 95%CI = 0.9274-0.9582 and 0.9564-0.9905, adj. p-value = 1.13×10^−12^ and, respectively). Our observation was consistent with the fact that the median age of IDH-mutants is younger than IDH-WT gliomas. Patients carrying the *IDH1* mutation generally had longer survival than those with IDH-WT^34^. Previous studies also reported that *IDH1* mutations often co-occurred with *ATRX* and *TP53* mutations, and mutations in these three genes were more prevalent in gliomas without *EGFR* mutations^15,35^. Indeed, we found that *EGFR* mutations were more common in older low-grade glioma patients (OR = 1.0865, 95%CI = 1.0525-1.1258, adj. p-value = 4.35×10^−7^) (Fig. 4f). Moreover, our SCNA analysis revealed an increase in the gain of *EGFR* with age in low-grade glioma but not in glioblastoma (Fig. 3d), suggesting the difference in age-associated genomic landscape between the two glioma types. Together with the SCNA results, gliomas from younger patients are associated with *IDH1*, *ATRX*, and *TP53* mutations, lower SCNAs, and longer survival. In contrast, gliomas from older patients were more likely to be IDH-WT with *EGFR* mutations, chromosome 7 gain and 10 loss, *CDKN2A* deletion and worse prognosis.

Mutations in cancer driver genes showed a positive or negative association with age depending on cancer types. For instance, *PTEN* mutations decreased with patient’s age in colon (OR = 0.9347, 95%CI = 0.8935-0.9738, adj. p-value = 0.0029) and endometrial cancers (OR = 0.9586, 95%CI = 0.9331-0.9840, adj. p-value = 0.0033) but increased with age in cervical cancer (OR = 1.0550, 95%CI = 1.0174-1.0959, adj. p-value = 0.0067). *CDH1* mutations were more frequent in younger stomach cancer patients (OR = 0.9414, 95%CI = 0.9027-0.9800, adj. p-value = 0.0061) but more common in older breast cancer patients (OR = 1.0218, 95%CI = 1.0049-1.0392, adj. p-value = 0.0171). These results highlight cancer-specific patterns of genomic alterations in relation to age. Overall, our results demonstrate that non-silent mutations in cancer driver genes were not uniformly distributed across ages and we have comprehensively identified, based on data available at present, genes that show age-associated mutation patterns. These patterns might point out age-associated disparities in carcinogenesis, molecular subtypes and survival outcome.

### Age-associated alterations in oncogenic signalling pathways

As we have identified numerous age-associated alterations in cancer driver genes in both SCNA and somatic mutation levels, we asked if the age-associated patterns also exist in particular oncogenic signalling pathways. We used the data from a previous TCGA study, which had comprehensively characterized 10 highly altered signalling pathways in cancers^36^. To make the subsequent analysis comparable to previous analyses, we restricted the analysis to samples that were used in our previous analyses, yielding 8,055 samples across 33 cancer types (Table 1). Using logistic regression adjusting for gender, race and cancer type, we identified five out of 10 signalling pathways that showed a positive association with age (adj. p-value < 0.05), indicating that the genes in these pathways are altered more frequently in older patients, concordant with the increase in overall mutations and SCNAs with age (Fig. 5a, Supplementary Table 11). The strongest association was found in cell cycle (OR = 1.0122, 95%CI = 1.0076-1.0168, adj. p-value = 1.40×10^−6^) and Wnt signalling (OR = 1.0122, 95%CI = 1.0073-1.0172, adj. p-value = 6.39×10^−6^). We next applied logistic regression to investigate the cancer-specific association between age and oncogenic signalling alterations for cancer types that contained at least 100 samples. In total, we identified 28 significant associations across 15 cancer types (adj. p-value < 0.05) (Fig. 5b, Supplementary Table 11). Alterations in Hippo and TP53 signalling pathways significantly associated with age, both positively and negatively, in five cancer types. Consistent with a pan-cancer analysis, cell cycle, Notch and Wnt signalling each showed an increase in alterations with age in three cancer types. We found that alterations in cell cycle pathway increased with age in low-grade glioma (OR = 1.0313, 95%CI = 1.0161-1.0467, adj. p-value = 0.00035). This was largely explained by the increase in *CDKN2A* and *CDKN2B* deletions with age as well as epigenetic silencing of *CDKN2A* in older patients (Fig. 5c). On the other hand, TP53 pathway alteration was more pronounced in younger patients (OR = 0.9520, 95%CI = 0.9372-0.9670, adj. p-value = 2.63×10^−8^), due to the mutations in the *TP53* gene (Fig. 5c). In endometrial cancer, two pathways – Hippo (OR = 0.9681, 95%CI = 0.9459-0.9908, adj. p-value = 0.0126) and Wnt (OR = 0.9741, 95%CI = 0.9541-0.9946, adj. p-value = 0.0240) - showed a negative association with age, that may be explained by the presence of hypermutated tumours in younger patients. Collectively, we reported pathway alterations in relation to age in several cancer types, highlighting differences in oncogenic pathways that might be important in cancer initiation and progression in an age-related manner.

**Fig. 5.**
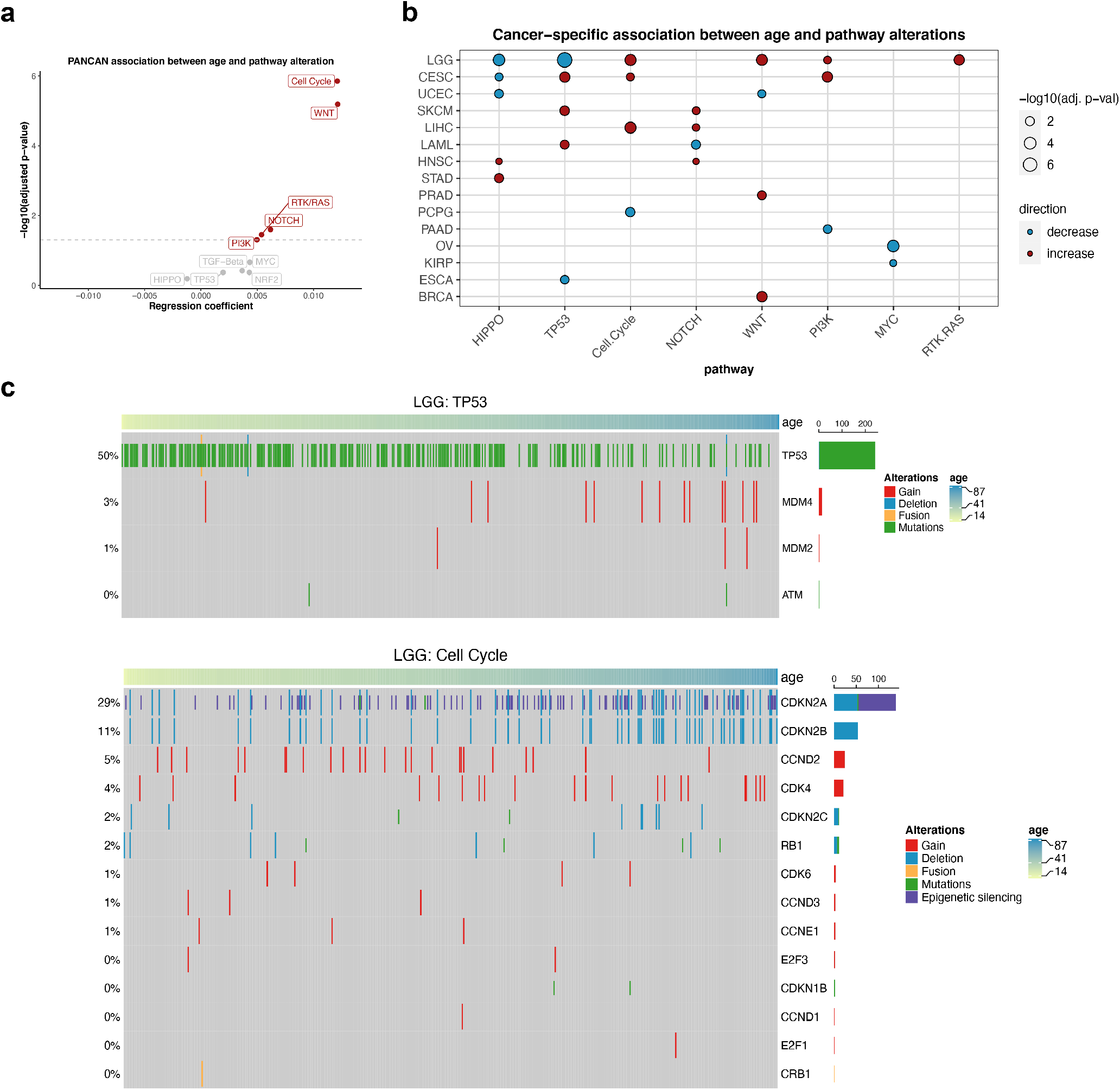
Association between cancer patients’ age and oncogenic signalling pathway alterations. **a** Association between age and oncogenic pathway alterations in the pan-cancer level. Multiple logistic regression coefficients and significant values are shown. Pathways with a significant positive association between age and alterations (adj. p-value < 0.05) are highlighted in red. **b** Cancer-specific age-associated pathway alterations. Pathways that show a significant positive and negative association with age per cancer type (multiple logistic regression, adj. p-value < 0.05) are displayed in red and blue dots, respectively. **c** Heatmap showing age-associated alterations in genes associated with TP53 and cell cycle pathways in LGG. Samples are sorted by age. Colours represent types of alteration.

### Age-associated gene expression and DNA methylation changes

Apart from the genomic differences with age, we investigated age-associated transcriptomic and epigenetic changes across cancers. We separately performed multiple linear regression analyses on gene expression data and methylation data of 24 cancer types that contained at least 100 samples in both types of data (Table 1). We noticed that, across all genes, the regression coefficient of age on gene expression negatively correlated with the regression coefficient of age on methylation in all cancer types (Supplementary Fig. 8), suggesting that the global changes of gene expression and methylation with age are in the opposite direction. This supports the established role of DNA methylation in suppressing gene expression. Numbers of significant differentially expressed genes with age (age-DEGs) (adj. p-value < 0.05, Supplementary Table 12) varied from nearly 5,000 up- and down-regulated genes in low-grade glioma to no significant gene in 5 cancers. Similarly, we also identified significant differentially methylated genes with age (age-DMGs, Supplementary Table 13) (adj. p-value < 0.05), the number of age-DEGs and age-DMGs were consistent for most cancer types (Fig. 6a). We next focused our analysis on 10 cancer types that contained at least 150 age-DEGs and 150 age-DMGs, including low-grade glioma, breast cancer, endometrial cancer, oesophageal cancer, papillary renal cell carcinoma, ovarian cancer, liver cancer, acute myeloid leukaemia, melanoma, and prostate cancer. We identified overlapping genes between age-DEGs and age-DMGs and found that most of them, from 84% (37/44 genes) in ovarian cancer to 100% in acute myeloid leukaemia (57 genes) and prostate cancer (7 genes), were genes that presented increased methylation and decreased expression with age and genes that had decreased methylation and increased expression with age (Fig. 6b-c, Supplementary Fig. 9, Supplementary Table 14). We further examined the correlation coefficient between methylation and expression comparing between 4 groups of genes 1) overlap genes between age-DMGs and age-DEGs (age-DMGs-DEGs), 2) age-DMGs only, 3) age-DEGs only, and 4) other genes. We found that age-DMGs-DEGs had the most negative correlation between DNA methylation and expression when comparing with other groups of genes (Fig. 6d, Supplementary Fig. 10, Supplementary Table 15), highlighting that age-associated gene expression changes in cancer are repressed, at least in part, by DNA methylation.

**Fig. 6.**
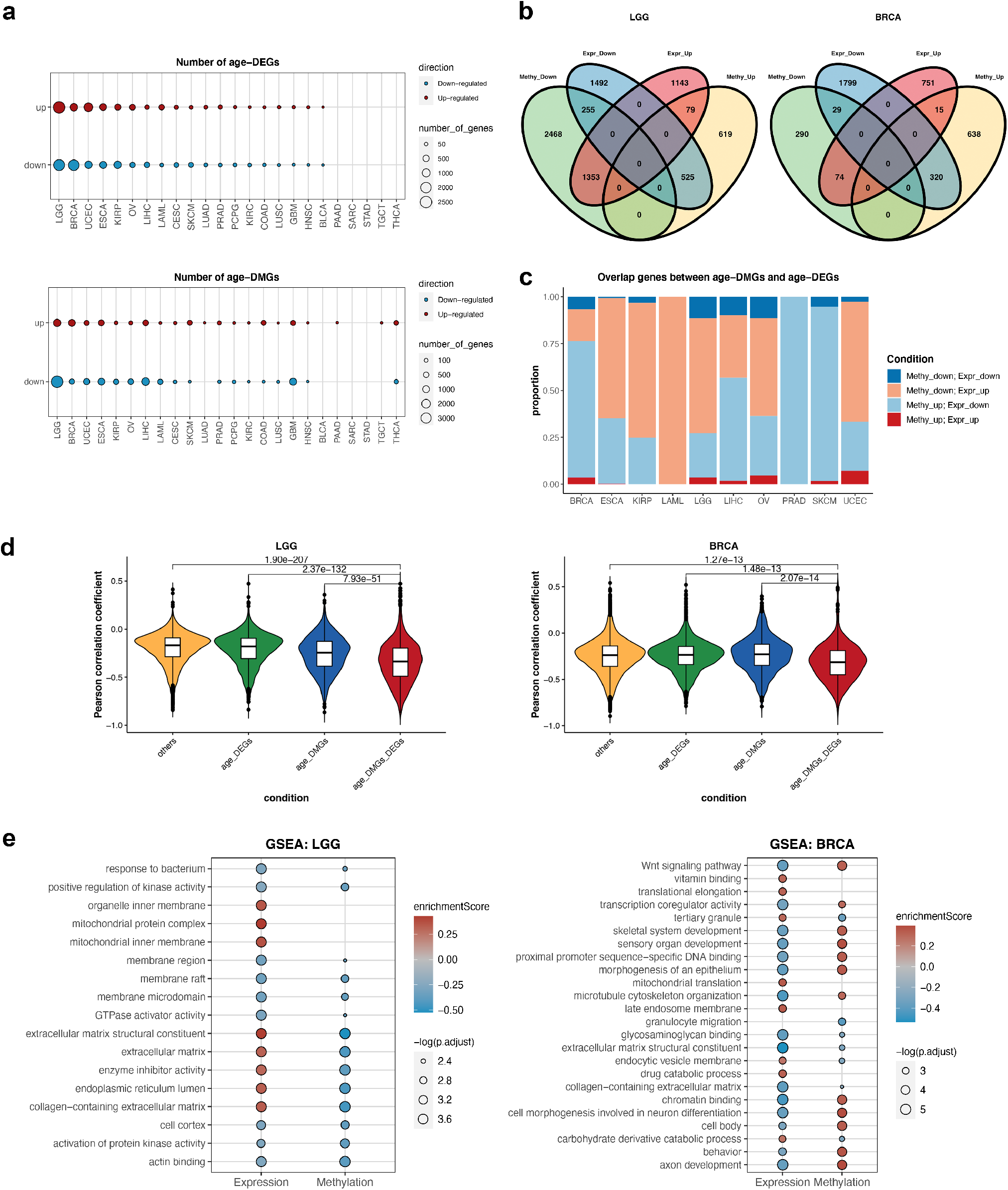
Age-related gene expression in cancers was controlled by age-related methylation. **a** Number of age-DEGs and age-DMGs across cancer types. Red dots represent up-regulated genes, while blue dots denote down-regulated genes. The dot size corresponds to the number of genes. **b** Venn diagrams of the overlap between age-DEGs and age-DMGs. LGG and BRCA are shown as examples. Venn diagrams of the other cancers are shown in Supplementary Fig. 9. **c** The distribution of overlap genes between age-DMGs and age-DEGs. The genes were classified into (1) down-regulated methylation and down-regulated expression, (2) down-regulated methylation and up-regulated expression, (3) up-regulated methylation and down-regulated expression, and (4) up-regulated methylation and up-regulated expression. **d** Violin plots showing the distribution of the Pearson correlation coefficient between methylation and gene expression in LGG and BRCA. Genes were grouped into (1) common genes between age-DMGs and age-DEGs (age-DMGs-DEGs), (2) age-DMGs only genes, (3) age-DEGs only genes, and (4) other genes. The group comparison was performed by the Kruskal-Wallis test. The pairwise comparisons were done using Dunn’s test. P-values from Dunn’s test between age-DMGs-DEGs and the other groups are shown. The plots for the other cancers are shown in Supplementary Fig. 10. **e** The enriched gene ontology (GO) terms identified by GSEA in LGG and BRCA. The dot size corresponds to a significant level. A GO term was considered significantly enriched term if adj. p-value < 0.05 for gene expression and adj. p-value < 0.1 for methylation. Colours represent enrichment scores, red denotes positive score (enriched in older patients), while blue signifies negative score (enriched in younger patients). The plots for the other cancers are shown in Supplementary Fig. 11.

We next performed Gene Set Enrichment Analysis (GSEA) to gain biological insights into the expression and methylation changes with age. We identified various significantly enriched Gene Ontology (GO) terms across cancers (Fig. 6e, Supplementary Fig. 11, Supplementary Table 16). Notably, several GO terms were enriched in both expression and methylation changes, in the opposite direction. The enriched terms in breast cancer included several signalling, metabolism, and developmental pathways. The Wnt signaling pathway, which was altered more frequently in older breast cancer patients (Fig. 5b), showed a decrease in gene expression and increase in methylation with age. In low-grade glioma, interestingly, mitochondrial terms were enriched in the gene expression of younger patients. Mitochondrial dysfunction is known to be important in glioma pathophysiology^37^, thus the different levels of mitochondrial aberrations might contribute to the disparities in the aggressiveness of gliomas in patients of different age. We also identified numerous immune-related terms enriched across several cancer types, including oesophageal, papillary renal cell, liver, and prostate cancers (Supplementary Fig. 11, Supplementary Table 16). Previous studies suggested alterations in immune-related gene expression and immune cell abundance changes with age in cancers^38,39^. In the present study, we have systematically characterised the transcriptome and methylation in relation to age across cancer types. Our results suggest that gene expression changes with age in cancer are controlled, at least in part, by DNA methylation. These changes reflect differences in biological pathways that might be important in tumour development.

## Discussion

Although age is an important risk factor for cancer, how age impacts the molecular landscape of cancer is not well understood. In this study, we provide a comprehensive overview of the age-associated molecular landscape in cancer, including genomic instability, LOH, WGD, SCNAs, somatic mutations, pathway alterations, gene expression, and DNA methylation. We confirmed the known increase in mutation load^4,5^ and found an increase in genomic instability, LOH and WGD with age in several cancer types. We identified several age-related pan-cancer and cancer-specific alterations. The highest age-related differences were evident in low-grade glioma and endometrial cancer.

Cancer develops through the accumulation of genetic and epigenetic alterations. Mutation accumulation with age is thought to be a cause of cancer and a substantial portion of mutations arise before cancer initiation^6^. The age-associated mutation accumulation has been demonstrated in both cancer^4,5^ and normal tissues^40–42^, providing a better understanding of an early carcinogenesis event. Our results show that, in addition to mutations, SCNAs, LOH and WGD increase with age in several cancers, in particular low-grade glioma, endometrial and ovarian cancers. Recent evidence suggests that SCNA burden is a prognostic factor associated with both recurrence and death^43^, thus, an increased SCNA level with age might relate to poor prognosis in the elderly.

The negative association between age and mutation in *IDH1* and *ATRX* in glioma points towards the difference of patient age at diagnosis between the *IDH*-mutant and *IDH*-WT subtypes. *IDH*-mutant tumours are observed in the majority of low-grade glioma and show favourable prognosis. *IDH*-WT low-grade gliomas, on the other hand, more resemble glioblastomas and have poorer survival. In glioblastoma, although *IDH*-mutants are a minority of tumours, they are also associated with younger age^44^. The present study together with others^34,45^, therefore indicates that glioma shows unique age-associated subtypes. However, more research is needed to understand how age influences the evolution of glioma subtypes.

Our results highlighted substantial age-associated differences in the genome of endometrial cancer. Younger endometrial tumours associate with a *POLE* and MSI-H subtypes, leading to an enrichment of hypermutated tumours, while tumours from older patients tend to harbour a higher SCNA level and lower mutation load. Previous studies have classified endometrial cancer into four subtypes: *POLE*, MSI-H, copy-number low and copy-number high subtypes. The *POLE* subtype and MSI-H subtype are dominated by the *POLE* and defective mismatch repair mutational signatures, respectively^33^. Conversely, the copy-number low and copy-number high subtypes had a dominant ageing-related mutational signature^31^. The *POLE* and MSI-H subtypes have a favourable prognosis, while the copy-number high subtype is associated with poor survival. Therefore, endometrial cancer from younger patients is associated with *POLE* mutations, mismatch repair defects, high mutation load and better survival outcomes. Older endometrial cancer, however, is related to extensive SCNAs and worse prognosis. Importantly, apart from low-grade glioma and endometrial cancer, we demonstrate that other cancer types also present an age-associated genomic landscape in cancer driver genes and oncogenic signalling pathways. These results highlight the impact of age on the molecular profile of cancer.

Having identified these age-related differences in the molecular landscapes of various cancers, the obvious question is what drives these differences. Accumulating evidence has underscored the importance of tissue environment changes with ageing in cancer initiation and progression^7,8,39,46^. We reason that tissue environment changes during ageing and might provide different selective advantages for tumours harbouring different molecular alterations in turn directing the tumours to different evolutionary routes. Therefore, cancer with different genomic alterations might thrive better in younger or older patients. Gene expression and epigenetic changes related to ageing have been studied and linked to cancer^8,38,47,48^. Here, we identified numerous age-associated gene expression and corresponding DNA methylation in a broad range of cancers. Indeed, age-DMGs-DEGs are those with the strongest negative correlation between methylation and expression when comparing with other groups, indicating that differentially expressed genes with age in cancer are partly regulated by methylation. Expression and methylation changes with age link to several biological processes, showing that cancer from patients with different ages present different phenotypes. We also noticed that cancer in female reproductive organs including breast, ovarian and endometrial cancers are among those with the highest number of age-DEGs and age-DMGs. These cancers tend to have a higher mass-normalised cancer incidence, which may reflect evolutionary trade-offs involving selective pressures related to reproduction^49^. The age-associated hormonal changes could also be responsible for this age-related expression differences in cancer^50^. The limitation of this analysis is that although we have already included tumour purity in our linear model, it is not possible to account for the different tumour-constituent cell proportions and thus fully exclude the influence from gene expression of non-tumor cells such as infiltrating immune cells^39^. Further studies are required to provide mechanistic understanding of the impact of an ageing microenvironment in shaping tumour evolution.

During the preparation of our manuscript, a study based on a similar concept has been released by Li et al^51^. In this work, Li et al. used TCGA and the recent pan-cancer analysis of whole genomes (PCAWG) data to study the age-associated genomic differences in cancer. Results from the two studies are consistent on several points. Firstly, both studies indicate the increase in mutations and SCNA levels with age. Next, despite using slightly different statistical cutoffs and models, several age-associated genomic features are identified by both studies, for example, the higher frequency of *IDH1* and *ATRX* mutations in younger glioma patients. Li et al. explored mutational timing and signatures, which suggested the possible underlying mechanisms for age-associated genomic differences. Our study, however, has also featured an age-related genomic profile in endometrial cancer. We have investigated cancer-specific associations between age and LOH, WGD and oncogenic signalling. Furthermore, we have analysed age-related global transcriptomic and DNA methylation changes. Our study are complementary with the Li et al. study, both studies thus serve as a foundation for understanding age-related differences and effects on the cancer molecular landscape and emphasise the importance of age in cancer genomic research that is particularly valuable in the clinical practice.

## Methods

### Data acquisition

Publicly available copy-number alteration seg files (nocnv_hg19.seg), normalized mRNA expression in RSEM (.rsem.genes.normalized_results TCGA files from the legacy archive, aligned to hg19), and clinical data (XML files) from TCGA were downloaded using *TCGAbiolinks* (version 2.14.1)^52^. The mutation annotation format (MAF) file was downloaded from the TCGA MC3 project^53^ (https://gdc.cancer.gov/about-data/publications/mc3-2017). The somatic alterations in 10 canonical oncogenic pathways across TCGA samples were obtained from a previous study by Sanchez-Vega et al^36^. The TCGA Illumina HumanMethylation450K array data (in β-values) was downloaded from Broad GDAC Firehose (http://gdac.broadinstitute.org/). The allele-specific copy number, tumour ploidy, tumour purity were estimated using ASCAT (version 2.4.2)^54^ on hg19 SNP6 arrays with penalty=70 as previously described^55,56^. We restricted our subsequent analyses to samples that have these profiles available. WGD duplication was determined using fraction of genome with LOH and ploidy information. Genomic instability (GI) scores have been computed as fraction of genomic regions that are not in 1+1 (for non WGD tumours) or 2+2 (for WGD tumours) statuses. For each data type and each cancer type, the summary of the numbers of TCGA samples included in the analysis, alongside clinical variable analysed are presented in the Supplementary Table 1.

### Statistical analysis and visualisation

Simple linear regression and multiple linear regression adjusting for clinical variables were performed using the *lm* function in R to access the relationship between age and continuous variables of interest. Simple logistic regression to investigate the association between age and binary response (e.g. mutation as 1 and wild-type as 0) and multiple logistic regression adjusting for covariates were carried out using the *glm* function in R. In pan-cancer analyses, gender, race and cancer type were variables included in the linear model. Clinical variables used in cancer-specific analyses included gender, race, pathologic stage, neoplasm histologic grade, smoking status, alcohol consumption and cancer-specific variables such as oestrogen receptor (ER) status in breast cancer. To avoid the potential detrimental effect caused by missing data, we retained only variables with missing data less than 10% of samples used in the somatic copy number alteration analysis (Supplementary Table 1). To account for the difference in the proportion of cancer cells in each tumour, tumour purity (cancer cell fraction) estimated from ASCAT was included in the linear model. When necessary, to avoid the separation problem that might occur due to the sparse-data bias^57^, *logistf* function from the *logistf* package (version 1.23)^58^ was used to perform multivariable logistic regression with Firth’s penalization^59^. Effect sizes from logistic regression analyses were reported as odds ratio per year and 95% confidence intervals. P-values from the analyses were accounted for multiple-hypothesis testing using Benjamini–Hochberg procedure^60^. Statistical significance was considered if adj. p-value < 0.05, unless specifically indicated otherwise.

All statistical analyses were carried out using R (version 3.6.3)^61^. Plots were generated using *ggplot2* (version 3.3.2)^62^, *ggrepel* (version 0.8.2)^63^, *ggpubr* (version 0.4.0)^64^, *ComplexHeatmap* (version 2.2.0)^65^, and *VennDiagram* (version 1.6.20)^66^.

### GI score analysis

GI score was calculated as a genome fraction (percent-based) that does not fit the estimated tumour ploidy, 2 for normal diploid, and 4 for tumours that have undergone the WGD process. Simple linear regression was performed to identify the association between age and GI score. For pan-cancer analysis, multiple linear regression was used to adjust for gender, race, and cancer type. For cancer-specific analysis, multiple linear regression accounting for clinical variables was conducted on the cancer types that had a significant association between age and GI score from the simple linear regression analysis (adj. p-value < 0.05). The complete set of results is presented in Supplementary Table 2.

### Percentage genomic LOH quantification and analysis

To quantify the percent genomic LOH for each tumour, we used allele-specific copy number profiles from ASCAT. X and Y chromosome regions were discarded from the analysis. The LOH segments were segments that harbour only one allele. The percent genomic LOH was defined as 100 times the total length of LOH regions / length of the genome.

Simple linear regression and multiple linear regression adjusting for gender, race, and cancer types were conducted to investigate the relationship between age and the percent genomic LOH in the pan-cancer analysis. For cancer-specific analysis, simple linear regression was performed followed by multiple linear regression accounting for clinical factors for cancers with a significant association in simple linear regression analysis (adj. p-value < 0.05). The complete set of results is in Supplementary Table 3.

### WGD analysis

WGD status for each tumour was obtained from fraction of genome with LOH and tumour ploidy. To investigate the association between age and WGD across the pan-cancer dataset, we performed simple logistic regression and multiple logistic regression correcting for gender, race, and cancer type. For cancer-specific analysis, simple logistic regression was performed to access the association between age and WGD on tumours from each cancer type. Cancer types with a significant association between age and WGD (adj. p-value < 0.05) were further subjected to the multiple logistic regression accounting for the clinical variables. The complete set of results is in Supplementary Table 4.

### List of known cancer driver genes

We compiled a list of known cancer driver genes from (1) the list of 243 COSMIC classic genes from COSMIC database version 91^67^ (downloaded on 1^st^ July 2020), (2) the list of 260 significantly mutated genes from Lawrence et al^68^, and (3) the list of 299 cancer driver genes from the TCGA Pan-Cancer study^69^. In total, we obtained 505 cancer genes and focused on the mutations and focal-level SCNAs on these genes in our study. The full list of cancer driver genes is available in Supplementary Table 8.

### Recurrent SCNA analysis

Recurrent arm-level and focal-level SCNAs of each cancer type were identified using GISTIC2.0^22^. Segmented files (nocnv_hg19.seg) from TCGA, marker file and CNV file, provided by GISTIC2.0, were used as input files. The parameters were set as follows: ‘-genegistic 1 -smallmem 1 -qvt 0.25 -ta 0.25 -td 0.25 -broad 1 -brlen 0.7 -conf 0.95 -armpeel 1 -savegene 1’. Based on these parameters, broad events were defined as the alterations happen in more than 70% of an arm. The log2 ratio thresholds for copy number gains and deletions were 0.25 and −0.25, respectively. The confidence level was set as 0.95 and the q-value was 0.25.

To investigate the association between age and arm-level SCNAs for each cancer type, simple logistic regression was performed for each chromosomal arm that was identified as recurrent SCNA in a cancer type. Only cancer types with more than 100 samples were included in this analysis (Table 1). Arms with a significant association (adj. p-value < 0.05) were further adjusted for clinical variables using multiple logistic regression. The complete set of results is in Supplementary Table 6. Similarly, simple and multiple logistic regression was conducted on the focal-level SCNAs for each cancer type. Regions that are not overlapped with centromeres or telomeres were removed from the analysis. The complete set of results is in Supplementary Table 7.

To confirm the impact of SCNAs on gene expression, we investigated the correlation between GISTIC2.0 score and RNA-seq based gene expression (log2(normalised RSEM + 1)) for tumours that have both types of data using Pearson correlation. The correlation was considered significant if the p-value corrected for multiple-hypothesis testing using the Benjamini-Hochberg procedure < 0.05. The complete set of results is in Supplementary Table 7.

### SCNA score quantification and analysis

Previous studies have developed the SCNA score representing the SCNA level of a tumour^12,23^. We applied the methods described by Yuan et al^12^ to calculate SCNA scores. Using SCNA profiles from GISTIC2.0 analysis, SCNA scores for each tumour were derived at three different levels (chromosome-, arm-, and focal-level). For each tumour, each focal-event log2 copy number ratio from GISTIC2.0 was classified into the following score: 2 if the log2 ratio ≥ 1, 1 if the log2 ratio < 1 and ≥ 0.25, 0 if the log2 ratio < 0.25 and ≥ −0.25, −1 if the log2 ratio < −0.25 and ≥ −1, and −2 if the log2 ratio < −1. The |score| from each focal event in a tumour was then summed into a focal score of a tumour. Thereafter, the rank-based normalisation (rank/number of tumours in a cancer type) was applied to focal scores from all tumours within the same cancer type, resulting in normalized focal-level SCNA scores. Therefore, tumours with high focal-level SCNAs will have focal-level SCNA scores close to 1, while tumours with low focal-level SCNAs will have scores close to 0. For the arm- and chromosome-level SCNA scores, a similar procedure was applied to the broad event log2 copy number ratio from GISTIC2.0. An event was considered as a chromosome-level if both arms have the same log2 ratio, otherwise it was considered as an arm-level. Similar to the focal-level SCNA score, each arm- and chromosome-event log2 copy number ratio was classified into the 2, 1, 0, −1, −2 scores using the threshold described above. The |score| from all arm-events and chromosome-events for a tumour were then summed into an arm score and chromosome score, respectively. For each cancer type, the rank-based normalisation was applied to arm scores and chromosome scores from all tumours to derive normalised arm-level SCNA scores and normalised chromosome-level SCNA scores, respectively. An overall SCNA score for a tumour was defined as the sum of focal-level, arm-level, and chromosome-level SCNA scores. A chromosome/arm-level SCNA score for a tumour was defined as the sum of chromosome-level and arm-level SCNA scores.

The association between age and overall, chromosome/arm-level, and focal-level SCNA scores for each cancer type was investigated using simple linear regression. Cancer types with a significant association (adj. p-value < 0.05) were then subjected to multiple linear regression analysis adjusting for the clinical variables. The complete set of results is included in Supplementary Table 5.

### Analysis of age-associated somatic mutation in cancer genes

We obtained the mutation data from the MAF file from the recent TCGA Multi-Center Mutation Calling in Multiple Cancers (MC3) project^53^. In the MC3 effort, variants were called using seven variant callers. We filtered the variants to keep only non-silent SNVs and indels located in gene bodies, retaining only “Frame_Shift_Del”, “Frame_Shift_Ins”, “In_Frame_Del”, “In_Frame_Ins”, “Missense_Mutation”, “Nonsense_Mutation”, “Nonstop_Mutation”, “Splice_Site” and Translation_Start_Site in the “Variant_Classification” column. We focused only on mutations in the cancer genes from our compiled list of cancer driver genes. To prevent the bias that might cause by hypermutated tumours, we restricted the analysis to tumours with < 1,000 mutations per exome. For pan-cancer analysis, multiple logistic regression accounting for gender, race and cancer type was performed to investigate the association between age and mutations in 20 cancer genes that are mutated in > 5% of samples (Supplementary Table 10). For cancer-specific analysis, simple logistic regression was used to identify cancer genes that the mutations in these genes are associated with the patient’s age. Only genes that are mutated in > 5% of samples from each cancer type were included in the analysis. The significant associations (adj. p-value < 0.05) were further investigated using multiple logistic regression accounting for clinical variables. The complete set of results is in Supplementary Table 10.

### Analysis of mutational burden, MSI-H status, and *POLE*/*POLD1* mutations

A mutational burden was defined as the total non-silent mutations in an exome. The mutational burden for each tumour was log-transformed before using it in the subsequent analysis. To investigate the relationship between age and mutational burden in pan-cancer, multiple linear regression adjusting for gender, race and cancer type was conducted. For cancer-specific analysis, simple linear regression was performed. Cancer types with a significant association between age and mutational burden in simple linear regression analysis (adj. p-value < 0.05) were further examined using multiple linear regression accounting for clinical factors. The complete set of results is in Supplementary Table 9.

Microsatellite instability status for COAD, READ, STAD, and UCEC were downloaded from TCGA using *TCGAbiolinks*. To study the association between the presence of high microsatellite instability (MSI-H) and age, tumours were divided into binary groups: MSI-H = TRUE and MSI-H = FALSE. Multiple logistic regression adjusting for clinical variables was then performed. Similarly, *POLE* and *POLD1* mutation status were in a binary outcome (mutated and not mutated). Multiple logistic regression was used to investigate the association between age and *POLE*/*POLD1* mutations in cancer types that contained POLE/POLD1 mutations in > 5% of samples.

### Oncogenic signalling pathway analysis

We used the list of pathway-level alterations in ten oncogenic pathways (cell cycle, Hippo, Myc, Notch, Nrf2, PI-3-Kinase/Akt, RTK-RAS, TGFβ signaling, p53 and β-catenin/Wnt) for TCGA tumours comprehensively complied by Sanchez-Vega et al^36^. Member genes in the pathways were accessed for SCNAs, mutations, epigenetic silencing through promoter DNA hypermethylation and gene fusions. We retained only the pathway alteration data of samples that were presented in our SCNA analysis. For the pan-cancer analysis, we employed multiple logistic regression adjusting for the patient’s gender, race and cancer type to demonstrate the relationship between pathway-level alteration and age. To investigate the association between age and cancer-specific pathway alterations, we performed simple logistic regression. Cancer types with a significant association (adj. p-value < 0.05) were further examined by multiple logistic regression accounting for clinical variables. The complete set of results is in Supplementary Table 11.

### Gene expression and DNA methylation analysis

To render the results from gene expression and DNA methylation comparable, we limited the analysis to genes that are presented in both types of data. The lowly expressed genes were filtered out from the analysis by keeping only genes with RSEM > 0 in more than 50 percent of samples. Only protein coding genes identified using biomaRt^70^ (Ensembl version 100, April 2020). Normalised mRNA expression in RSEM for each TCGA cancer type was log2-transformed before subjected to the multiple linear regression analysis adjusting for clinical factors. RNA-seq data for colon cancer and endometrial cancer consisted of two platforms, Illumina HiSeq and Illumina GA. Thus, a platform was included as another covariate in the linear regression model for these two cancer types. Genes with adj. p-value < 0.05 were considered significantly differentially expressed genes with age (age-DEGs) (Supplementary Table 12). DNA methylation data was presented as β-values, which are the ratio of the intensities of methylated and unmethylated alleles. Because multiple methylation probes can be mapped to the same gene, we used the one-to-one mapping genes and probes by selecting the probes that are most negatively correlated with the corresponding gene expression from the files meth.by_min_expr_corr.data.txt. Similar multiple linear regression to the gene expression analysis was performed on the methylation data. Genes with adj. p-value < 0.05 were considered significant differentially methylated genes with age (age-DMGs). The complete set of results is in Supplementary Table 13.

The correlation between gene expression and DNA methylation was calculated using Pearson correlation. We used the Kruskal-Wallis test to investigate the differences between correlation coefficients among groups (age-DMGs-DEGs, age-DMGs, age-DEGs, other genes). The pairwise comparisons were carried out by Dunn’s test. The complete set of results is in Supplementary Table 15.

Gene Set Enrichment Analysis (GSEA) was performed to investigate the Gene Ontology (GO) terms that are enriched in tumours from younger or older patients. The analysis was done using the package *ClusterProfiler* (version 3.14.3)^71^. The complete list of enriched GO terms is presented in Supplementary Table 16.

## Data availability

TCGA data used in this study are publicly available and can be obtained from NCI’s Genomic Data Commons portal (https://portal.gdc.cancer.gov/), *TCGAbiolinks* (version 2.14.1)^52^ and Broad GDAC Firehose (http://gdac.broadinstitute.org/).

## Code availability

The custom scripts for data analysis and generate figures are available at https://github.com/maglab/Age-associated_cancer_genome.

## Acknowledgements

K.C. is supported by a Mahidol‐Liverpool PhD scholarship from Mahidol University, Thailand, and the University of Liverpool, UK. J.P.M. is grateful to funding from the Wellcome Trust (208375/Z/17/Z) and the Biotechnology and Biological Sciences Research Council (BB/R014949/1). This work was supported by the Francis Crick Institute, which receives its core funding from Cancer Research UK (FC001202), the UK Medical Research Council (FC001202), and the Wellcome Trust (FC001202). P.V.L. is a Winton Group Leader in recognition of the Winton Charitable Foundation’s support towards the establishment of The Francis Crick Institute. We wish to thank members of the Integrative Genomics of Ageing Group for suggestions and discussion.

## Author contributions

K.C., T.L., P.V.L. and J.P.M. conceived the project and designed the study. T.L. and P.V.L. provided data. K.C. performed the analyses with helps from T.L. T.L., L.P., P.V.L. and J.P.M. provided critical insights and were involved in data interpretation. K.C. wrote the first draft of the manuscript. All authors edited and approved the manuscript.

---

**Supplementary Fig. 1** Association between age and (**a**) GI score and (**b**) percent genomic LOH. Multiple linear regression was performed to identify the relationship between age and GI score or percent genomic LOH for each cancer type. Cancer types with a significant association (adj. p-value < 0.05) are shown together with adjusted R-squared and p-values from multiple linear regression analysis.

**Supplementary Fig. 2** Association between age and (**a**) overall SCNA score, (**b**) chromosome/arm-level SCNA score, and (**c**) focal-level SCNA score. Multiple linear regression was performed to identify the relationship between age and SCNA score for each cancer type. Cancer types with a significant association (adj. p-value < 0.05) are shown together with adjusted R-squared and p-value from multiple linear regression analysis.

**Supplementary Fig. 3** Heatmaps represent recurrent gain and deletion of arms across cancer types. Samples are sorted by age. Colours represent copy-number changes from GISTIC2.0, blue denotes deletion and red corresponds to gain.

**Supplementary Fig. 4** Heatmaps represent recurrent gain and deletion of focal-regions that showed the age-associated patterns across cancer types. Samples are sorted by age. Colours represent copy-number changes from GISTIC2.0, blue denotes deletion and red corresponds to gain. The direction legend shows whether the gain/deletion of the region increases or decreases with age.

**Supplementary Fig. 5** Association between age and mutational burden. Simple linear regression was performed to investigate the association between age and mutational burden. Cancer types with a significant association (adj. p-value < 0.05) were further investigated using multiple linear regression. The figure shows pan-cancer analysis and cancer type-specific analysis for every cancer that had a significant association in the simple linear regression analysis. Adjusted R-squared and p-value from multiple linear regression analysis are displayed.

**Supplementary Fig. 6** Association between age and (**a**) MSI-H in COAD, READ, and STAD, and (**b**) *POLE*/*POLD1* mutations in cancer types containing the mutations in these genes in more than 5% of the samples. The statistical significance (p-value showed) was calculated from the multiple logistic regression adjusting for clinical variables.

**Supplementary Fig. 7** Heatmap showing age-associated mutations in 11 cancer types. Samples are sorted by age. Colours represent types of mutation. The right annotation legend indicates the direction of change, increase or decrease mutations with age. The mutational burden of samples is presented in the dot above the heatmap.

**Supplementary Fig. 8** Pearson correlation between linear regression coefficient of age on DNA methylation level and linear regression coefficient of age on gene expression. The regression coefficients were obtained from the multiple linear regression analysis to investigate the association between age and DNA methylation or gene expression. Pearson correlation coefficient and p-values are shown.

**Supplementary Fig. 9** Venn diagrams of the overlap between age-DEGs and age-DMGs in 8 cancer types. Age-DEGs were separated into genes up-regulated with age (Expr_Up) and genes down-regulated with age (Expr_Down). Age-DMGs were classified into genes with increased methylation with age (Methy_Up) and genes with decreased methylation with age (Methy_Down).

**Supplementary Fig. 10** Pearson correlation coefficient between methylation and gene expression. **a** Violin plots showing the distribution of the Pearson correlation coefficient between methylation and gene expression in 8 cancer types. Genes were grouped into (1) overlapping genes between age-DMGs and age-DEGs (age-DMGs-DEGs), (2) age-DMGs only genes, (3) age-DEGs only genes, and (4) other genes. The group comparison was performed by the Kruskal-Wallis test. The pairwise comparisons were done using Dunn’s test. P-values from Dunn’s test between age-DMGs-DEGs and the other groups are shown. **b** Density plots showing the distribution of the Pearson correlation coefficient between methylation and gene expression across 10 cancer types.

**Supplementary Fig. 11** The enriched age-related gene ontology (GO) terms identified by GSEA in 8 cancer types. The dot size corresponds to a significant level. A GO term was considered significantly enriched if adj. p-value < 0.05 for gene expression and adj. p-value < 0.1 for methylation. Colours represent enrichment scores, red denotes positive score (enriched in older patients), while blue signifies negative score (enriched in younger patients). No enriched term was identified from the gene expression data of UCEC.

## List of supplementary tables

**Supplementary Table 1:** Summary of the number of samples and clinical variables used in the study

**Supplementary Table 2:** Association between age and GI scores

**Supplementary Table 3:** Association between age and percent genomic LOH

**Supplementary Table 4:** Association between age and WGD

**Supplementary Table 5:** Association between age and SCNA scores

**Supplementary Table 6:** Association between age and arm-level SCNAs

**Supplementary Table 7:** Association between age and focal-level SCNAs

**Supplementary Table 8:** List of previously identified cancer driver genes

**Supplementary Table 9:** Association between age and mutational burden

**Supplementary Table 10:** Association between age and somatic mutations

**Supplementary Table 11:** Association between age and oncogenic signalling pathway

**Supplementary Table 12:** Gene expression changes with age

**Supplementary Table 13:** DNA methylation changes with age

**Supplementary Table 14:** Number of overlapping genes between age-DEGs and age-DMGs

**Supplementary Table 15:** Comparision of the correlation between methylation and gene expression in age-DMGs-DEGs, age-DMGs, age-DEGs and other genes

**Supplementary Table 16:** List of enriched GO terms identified using GSEA

